# An Engineered IGF2 Mutant for Lysosomal Targeting Chimeras Development and Membrane Proteins Degradation

**DOI:** 10.1101/2024.02.20.581320

**Authors:** Yanchao Pan, Qing Xiang, Kai Deng, Muhammad Anwar, Leiming Wang, Yuan Wang, Qiulian Liang, Lirou Shen, Jing Yang, Weijun Shen

## Abstract

Lysosome-targeting chimeras (LYTACs) have emerged as a promising strategy for targeted degradation of membrane proteins, offering potential applications in drug development. Currently, two main methods for developing LYTACs exist: chemically modified antibodies ^[1-2]^ and wild-type insulin-like growth factor 2 (IGF2) fusion proteins (iLYTACs) ^[3]^. However, the fusion of the IGF2 arm within iLYTACs carries the risk of activating IGF1R tyrosine kinase activity and promoting tumor development. To address this concern, we introduce eiLYTACs, a technology that employs engineered IGF2 fusion antibodies to induce degradation of endogenous membrane proteins. Compared to the wild-type IGF2, the engineered IGF2 mutant exhibited minimal binding affinity for IGF1R but demonstrated a significant 100-fold increase in its binding affinity for IGF2R. In contrast to wild-type IGF2, which promotes tumor growth, the cells incubated with the engineered IGF2 showed no stimulation of tumor growth. The eiLYTACs strategy effectively inhibits tumor cell proliferation by degrading specific targets, resulting in a significant reduction in xenograft tumor size in experimental nude mice. More interestingly, our research revealed that eiLYTACs simultaneously degrade both homo- and heterodimers of disease-relevant proteins,which offer a promising strategy to address the activation of compensatory bypass signaling pathways, drug resistance, and tumor heterogeneity.

## 1. Introduction

Targeted protein degradation (TPD) has emerged as an effective strategy for treating cancer by selectively degrading oncogenic proteins that were previously considered challenging. Degrader technologies, such as proteolysis-targeting chimeras (PROTACs) ^[4]^ and molecular glues ^[5]^, exploit the ubiquitin proteasome system to facilitate targeted protein degradation. These approaches have demonstrated significant promise and have advanced into clinical trials ^[6]^. Lysosome-targeting chimeras (LYTACs), originally introduced by Bertozzi et al., represent another approach that utilizes ingeniously designed IgG-glycan bioconjugates to harness natural lysosome shuttling receptors such as ASGPR (asialoglycoprotein receptor) ^[2]^ or IGF2R (insulin-like growth factor 2 receptor) ^[1]^ for precise protein degradation. By utilizing these bioconjugates, LYTACs can specifically bind to these receptors on the cell surface, triggering internalization and subsequent trafficking of the target protein to the lysosomes for degradation. In addition to LYTACs, a diverse range of heterobifunctional molecules, including AbTACs ^[7]^, GlueTACs ^[8]^, KineTACs ^[9]^ and PROTABs ^[10]^, have been developed to form ternary complexes with a target protein, a lysosome trafficking receptor, or an E3 ligase, leading to the targeted degradation of membrane proteins. Unlike PROTACs, which are limited to targeting proteins with ligandable intracellular domains, large molecule-based degradation technologies like LYTACs can degrade extracellular and membrane-bound proteins. These proteins represent more than 40% of all protein-encoding genes and play crucial roles in various physiological and pathological processes, including cancer, age-related diseases, and autoimmune diseases ^[11]^. Recently, Zhang et al. reported another degradation approach called iLYTACs, which involves the genetic fusion of insulin-like growth factor 2 (IGF2) to either a nanobody or the IgG-binding Z domain ^[3]^.

Despite the significant progress made in the field of protein degradation, there are still certain challenges that need to be addressed. LYTACs have demonstrated effective degradation of membrane proteins. However, their preparation process is complex due to the need for chemical modifications. This complexity can pose challenges in terms of scalability and accessibility for LYTAC production. On the other hand, the iLYTAC technique offers the advantage of not requiring glycosylation modification, simplifying the overall process. However, it is important to note that the wild-type insulin-like growth factor 2 (IGF2) arm used in iLYTACs exhibits a high affinity for the type I IGF receptor (IGF1R). This high binding affinity raises concerns, as wild-type IGF2 is known to promote tumor growth through activation and signaling via IGF1R ^[12-13]^. In the context of tumors, the overexpression of IGF2 can further stimulate tumor growth by activating IGF1R signaling ^[14]^. To mitigate these potential side effects associated with wild-type IGF2, it is necessary to develop an engineered IGF2 variant that selectively binds to IGF2R with high potency while minimizing its binding to IGF1R.

In this study, we report the engineered IGF2-based fusion antibody strategy termed engineered IGF2 receptor-targeting chimeras (eiLYTACs), for targeted membrane protein degradation with high selectivity, specificity, and efficiency. Compared with the wild-type IGF2, the engineered IGF2-M5.6 mutant showed a 100-fold increase in binding affinity to IGF2R (Kd∼0.4nM). More importantly, IGF2-M5.6 mutant showed much reduced binding affinity towards IGF1R. This distinction is of great significance, as our experimental findings revealed that wild-type IGF2 has the potential to promote tumor cell growth. However, when cells were treated with the engineered IGF2-M5.6 mutant, no such tumor-promoting effects were observed. This indicates that the engineered IGF2-M5.6 mutant possesses improved selectivity and reduced interaction with IGF1R, mitigating the potential risks associated with IGF2-induced tumor growth. Our findings present evidence supporting the efficacy of engineered IGF2-bearing eiLYTACs in inducing rapid and extensive degradation of therapeutically relevant proteins in diverse cell types. Notably, our results demonstrate a unique feature of eiLYTACs: the ability to simultaneously degrade both homo- and heterodimers of disease-relevant proteins. By effectively degrading both types of protein complexes, eiLYTACs offer a versatile approach for addressing diseases where aberrant protein interactions play a crucial role. The ability to target and eliminate both homo- and heterodimers expands the scope of potential applications for eiLYTACs in treating various diseases and disorders.

## 2. Results and Discussion

### 2.1 Engineering of Insulin-like Growth Factor 2 (IGF2) for Selective IGF2R Binding and Targeted Protein Degradation

Previous studies have established that wild-type IGF2 can bind to IGF1R with high affinity^[12]^ and promote IGF1R autophosphorylation and signaling ^[15-16]^. This raises concerns about the potential promotion of tumor cell growth. We incubated SK-BR-3 cells with varying concentrations of hFc-IGF2 fusion protein and continuously monitored cell growth by IncuCyte (Sartorius, Incucyte S3). The results clearly demonstrated that hFc-IGF2 significantly stimulated the growth of SK-BR-3 cells (**Figure** 1), consistent with previous studies indicating that wild-type IGF2 promotes proliferation of breast cancer cells ^[17]^. To validate our findings, we further incubated hFc-IGF2 with BT474 breast cancer cells and SK-Hep1 hepatocellular carcinoma cells. Remarkably, we observed a substantial enhancement of proliferation in both cell types (Figure S1). This observation suggests that wild-type IGF2 can promote tumor proliferation in various types of cancer cells, complicating its potential application in membrane protein degradation as anti-tumor therapy.

**Figure 1.**
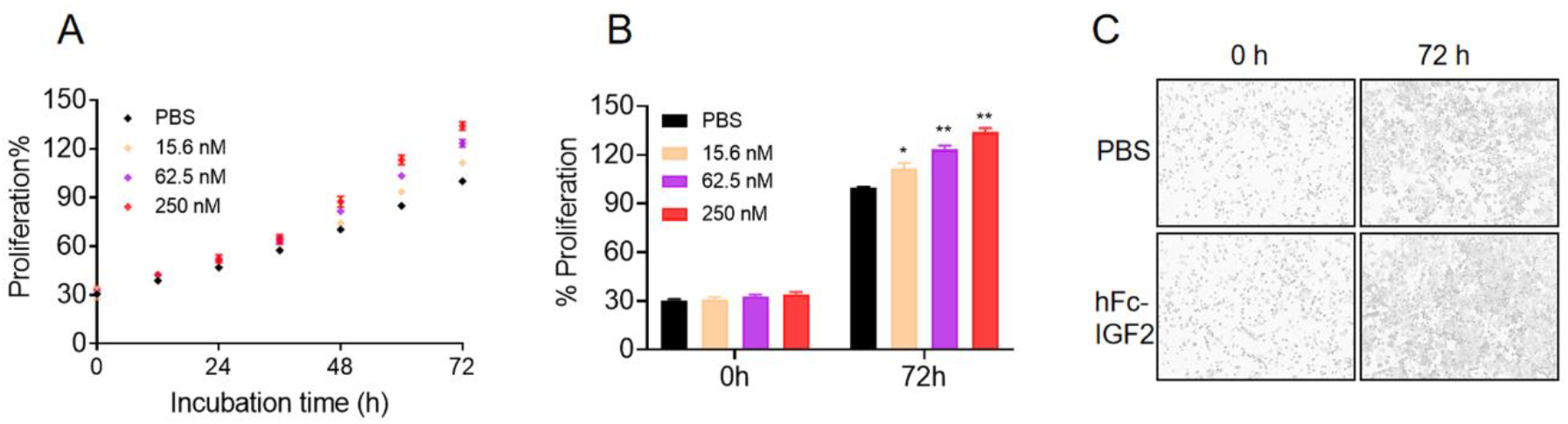
Promotion of SK-BR-3 cell proliferation by hFc-IGF2. A) Percentage of SK-BR-3 cell proliferation over a 3-day period. SK-BR-3 cells were treated with concentrations ranging from 15.6 to 250 nM of hFc-IGF2 on day 0. B) Statistical data on cell proliferation on the first and third days. C) Live imaging of SK-BR-3 cells using IncuCyte throughout the 3-day period following treatment with 250nM of hFc-IGF2.

To address the tumor promoting issue of wild-type IGF2, we decide to evolve wild-type IGF2 to obtain IGF2 mutant that can selectively and efficiently bind to IGF2R over IGF1R. Based on the crystal structure of IGF2/IGF2R ^[14, 18-19]^, we designed different IGF2 mutants and conducted two rounds of screening. Firstly, six IGF2 mutants coding gene were synthesized and constructed to the C-terminus of hFc with a (G)_5_ linker, using overlapping extension PCR and the synthetic oligonucleotide primers. These hFc-IGF2 mutants were tested for their binding affinity towards IGF1R and IGF2R. Our data revealed that the IGF2-M5 mutant, containing E6R, R37A, and V43M mutations, exhibited significantly enhanced affinity for IGF2R (Figure S2A). Subsequently, we fused IGF2-M5 to the C-terminus of the heavy chain of pertuzumab, an FDA-approved anti-HER2 antibody, and evaluated its impact on SK-BR-3 cell proliferation. Encouragingly, the fusion of IGF2-M5 to pertuzumab’s heavy chain resulted in a significant inhibition of SK-BR-3 cell proliferation, unlike the fusion of wild-type IGF2, which promoted cell growth (Figure S2B to S2D). These results demonstrate the potential of the evolved IGF2 mutant, IGF2-M5, to effectively inhibit tumor cell growth when fused to pertuzumab. Based on these results, we designed an additional set of ten IGF2 mutants for the second round of screening. These mutants were fused to the C-terminus of hFc and subjected to candidate screening via ELISA. Our analysis identified four IGF2 mutants with significantly enhanced binding affinity for IGF2R. Among these mutants, IGF2-M5.6, harboring E6R, F19L, R37A, and V43M mutations, exhibited the highest affinity for IGF2R, with a Kd value of approximately 0.4 Nm (**Figure** 2A). In addition, IGF2-M5.6 also demonstrated a remarkable decrease in binding affinity for IGF1R (Figure 2B). More importantly, no promotion of cell growth was observed in either breast cancer cells or HCC cells incubated with hFc-IGF2-M5.6 (Figures 2C, 2D and Figure S3). These findings underscore the potential of IGF2-M5.6 as a reliable candidate for targeted protein degradation. To demonstrate proof-of-concept application in eiLYTACs, we constructed HER2 targeting pertuzumab eiLYTAC (Ptz-eiLYTAC) by fusion of IGF2-M5.6 to the C-terminus of heavy chain of HER2 antibody pertuzumab with a (GGGGS)_3_ linker and co-expressed with pertuzumab light chain. We determined whether IGF2-M5.6 can trigger internalization of extracellular targets via IGF2R in a HER2 positive breast cancer cell line SK-BR-3, with pertuzumab as the control. Briefly, SK-BR-3 cells were incubated with Ptz-eiLYTAC together with goat anti-human IgG-647 for 1 h, and intracellular fluorescence was analyzed by confocal microscopy. The results demonstrated a high signal of IgG-647 within the cells, which colocalized with lysotracker, indicating that Ptz-eiLYTAC trafficked to the lysosomes (Figure 2E). In contrast, no internalization of IgG-647 was observed in the control group. Taken together, these data suggests that the engineered IGF2-M5.6 mutants can selectively bind to IGF2R which efficiently trigger internalization of extracellular targets.

**Figure 2.**
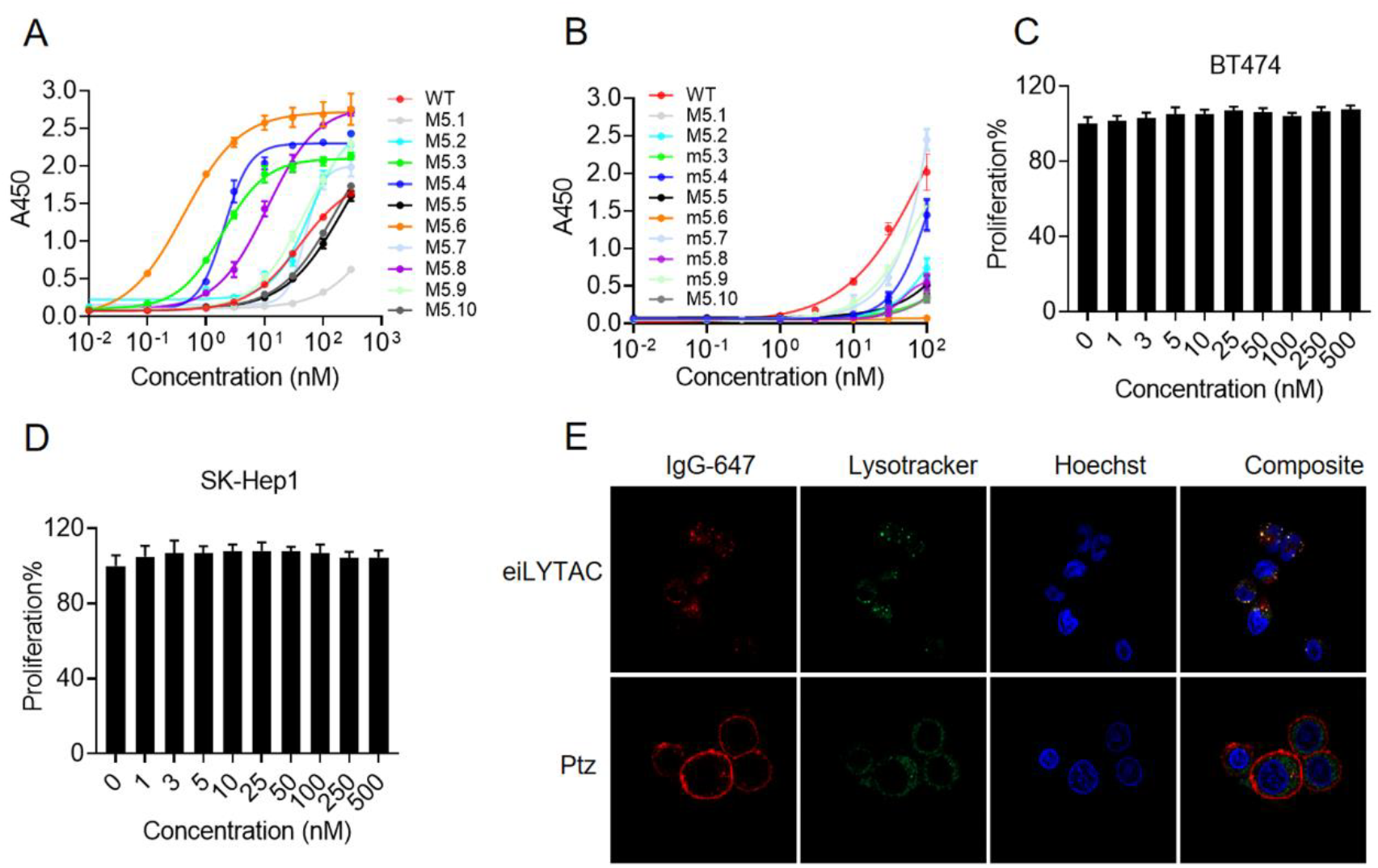
IGF2-M5.6 selectively binds to IGF2R and triggers efficient internalization of extracellular IgG-647. A) IGF2R binding properties of IGF2 analogues. Immunocaptured IGF2R was incubated with increasing concentrations of the following IGF2 analogues: wild-type IGF2 (WT) and IGF2 mutants (M5.1 to M5.10). B) IGF1R binding properties of IGF2 analogues. Immunocaptured IGF1R was incubated with increasing concentrations of the different IGF2 analogues listed above. C) and D) Proliferation of BT-474 and SK-Hep1 cells incubated with different concentrations of hFc-IGF2-M5.6 for 5 days. Cell proliferation was detected by CellTiter-Glo assay (Promega). E) Visualization of Ptz-eiLYTAC-mediated IgG-647 internalization in SK-BR-3 cells by confocal microscopy. The cells were incubated at 37 °C for 1 h with 10 nM pertuzumab (Ptz) or Ptz-eiLYTAC and 20 nM of Alexa Fluor 647-labeled goat anti-human secondary antibody in complete growth media. Cells were further labeled with LysoTracker Green for 30 min and Hoechst for 5 min before imaging.

### 2.2 EiLYTACs Degrade the Membrane Proteins and Induce Antiproliferative Effects

In the evaluation of eiLYTACs’ ability to degrade membrane proteins, we conducted experiments using SK-BR-3 cells and measured the degradation of HER2. The cells were incubated with varying concentrations of Ptz-eiLYTAC, and the degradation of HER2 was assessed through confocal microscopy and Western blot analysis. The results, depicted in **Figure** 3A and 3B, clearly demonstrated that the incubation of Ptz-eiLYTAC with SK-BR-3 cells effectively degraded the cell surface HER2. In contrast, cells incubated with pertuzumab showed no HER2 degradation. Consistently, degradation of HER2 was also observed in another two breast cancer cell lines (SUM159PT and MDA-MB-435), in a dose dependent manner. However, no degradation of HER2 was observed in cells treated with the pertuzumab even going up to 400 nM concentration (Figure S4A and S4B). Following the successful demonstration of eiLYTAC’s ability to induce lysosomal degradation of membrane proteins, we further investigated their anti-proliferation activity in cancer cells, comparing it to that of the parent antibody. SK-BR-3 breast cancer cells were treated with Ptz-eiLYTAC, and the proliferation of the cells was monitored over time using IncuCyte. As depicted in Figure 3C and 3D, Ptz-eiLYTAC decreased cell viability in SK-BR-3 cells in a concentration-dependent manner and much more potent than pertuzumab treatment alone.

**Figure 3.**
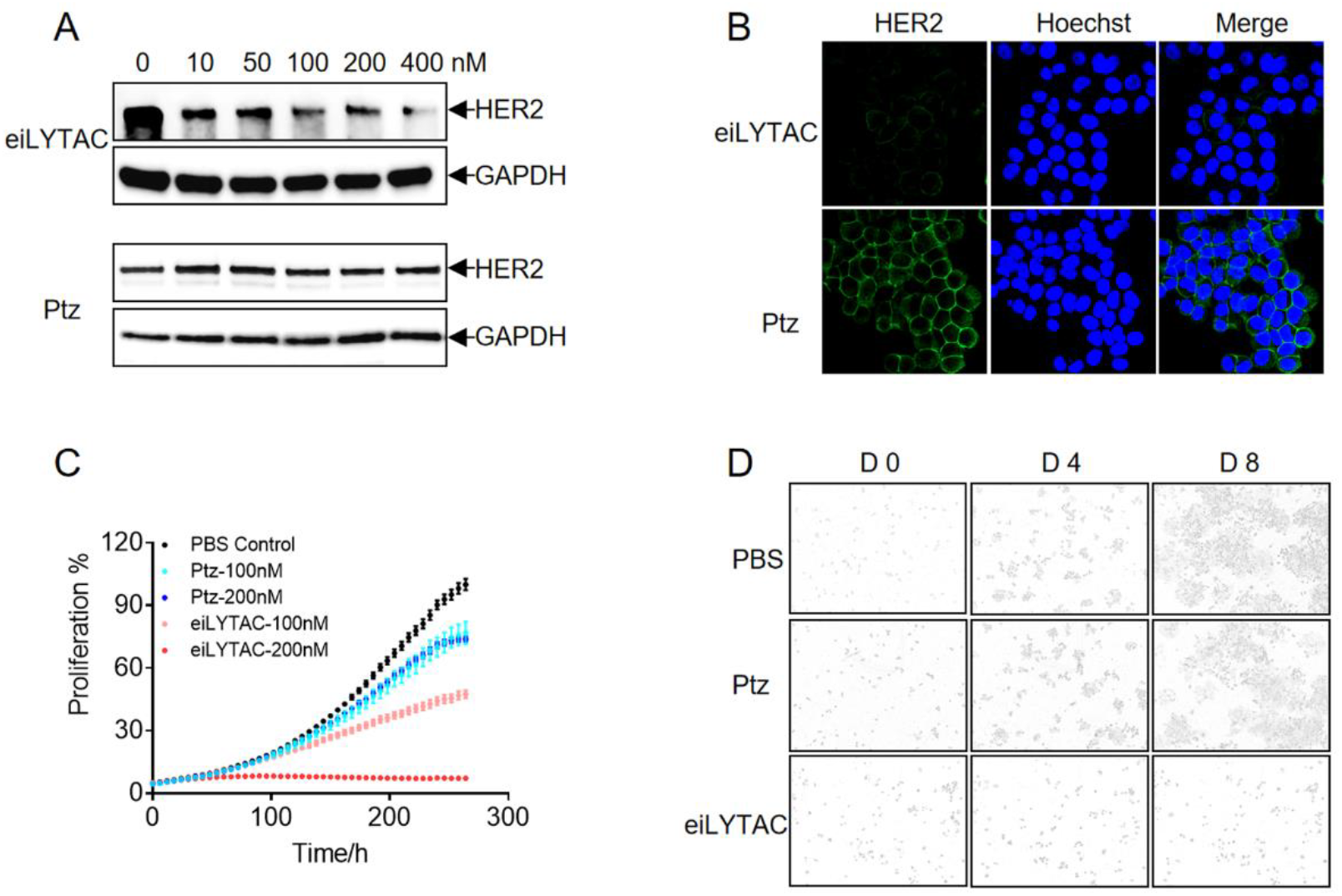
Ptz-eiLYTAC degrades the membrane protein HER2 and induces an antiproliferative effect in SK-BR-3 cells. A) Western blot analysis showing HER2 degradation in SK-BR-3 breast cancer cells following incubation with 0-400 nM of Ptz or Ptz-eiLYTAC for 48 h. B) Visualization of HER2 degradation in SK-BR-3 cells by confocal microscopy after treatment with 100 nM Ptz or Ptz-eiLYTAC for 48 h. C) Time course of percent proliferation of SK-BR-3 cells during a 264-hour treatment with 100 nM or 200 nM of Ptz or Ptz-eiLYTAC. D) Live imaging of SK-BR-3 cells using IncuCyte throughout at day 4 and day 8 following treatment with 100 nM or 200 nM of Ptz or Ptz-eiLYTAC.

To further validate the universality of eiLYTACs, we examined EGFR degradation in non-small cell lung cancer. The extensive mutational landscape of EGFR in certain cancers, such as non-small cell lung cancer, presents a challenge for targeted therapies that primarily focus on inhibiting the tyrosine kinase activity of EGFR. While sensitizing mutations in the tyrosine kinase domain have been effectively targeted by EGFR tyrosine kinase inhibitors (TKIs) like erlotinib, afatinib, and osimertinib, the emergence of additional mutations, including the C797S mutation, has been observed as a mechanism of acquired resistance to third-generation ^[20]^. Considering the extensive mutational landscape of EGFR and the need for therapies that can overcome new mutations in the tyrosine kinase domain ^[21]^, targeting the extracellular domain of EGFR with degraders presents a promising approach. To evaluate the efficacy of eiLYTACs in degrading EGFR, we generated Cet-eiLYTAC by fusing IGF2-M5.6 to the C-terminus of the heavy chain of cetuximab. The degradation efficiency of Cet-eiLYTAC was assessed in NCI-H1975 cell lines, which harbor the L858R and T790M mutation in exon 20 that confers resistance to first-generation tyrosine kinase inhibitors ^[22]^. As demonstrated in Figure S5A, Cet-eiLYTAC efficiently degraded EGFR, with over 90% of EGFR being degraded when the concentration of Cet-eiLYTAC reached 50 nM. Immunofluorescence staining of membrane-bound EGFR confirmed the degradation of EGFR in Cet-eiLYTAC-treated NCI-H1975 cells (Figure S5B). Considering that papulopustular rash (acne-like rash) develops in 80-86% of patients receiving cetuximab treatment ^[23]^, we designed another EGFR-targeting Nim-eiLYTAC by fusing IGF2-M5.6 to the C-terminus of the heavy chain of nimotuzumab. Nimotuzumab exhibits similar preclinical and clinical activity in certain indications but does not induce severe skin toxicity or severe hypomagnesemia or gastrointestinal adverse events ^[24]^. The degradation activity of Nim-eiLYTAC was evaluated in NCI-H1975 cells harboring the C797S mutation, which is known to confer resistance to the third-generation tyrosine kinase inhibitor osimertinib by impairing osimertinib’s covalent binding to the ATP-binding pocket of EGFR ^[25]^. Western blot analysis revealed that Nim-eiLYTAC efficiently degraded EGFR with C797S mutation (Figure S5C). Furthermore, Nim-eiLYTAC was demonstrated to inhibit NCI-H1975 cell growth (Figure S5D). Collectively, these results suggest that eiLYTACs exhibit broad-spectrum degradation preclinical activity against EGFR mutations.

### 2.3 EiLYTACs Degrade both Homo- and Heterodimers of Membrane Proteins

Despite the encouraging clinical results of trastuzumab, trastuzumab resistance remains a significant challenge in HER2-overexpressing breast cancers ^[26]^. *In vitro* studies have shown that HER2-positive JIMT-1 human breast cancer cells exhibit intrinsic resistance to trastuzumab due to epitope masking caused by extracellular matrix (ECM) components such as MUC4 and CD44-bound hyaluronic acid ^[27]^. Trastuzumab and pertuzumab target different epitopes on HER2, with trastuzumab binding to HER2 domain 4 proximal to the cell membrane and pertuzumab binding to domain 2 in the dimerization arm distal from the membrane ^[28-29]^. It is possible that pertuzumab can still bind to HER2 in tumors where ECM components interfere with trastuzumab binding. Based on this concept, we hypothesize that PTZ-eiLYTAC has the potential to overcome trastuzumab resistance caused by epitope masking. To test this hypothesis, we treated JIMT-1 cells with varying concentrations of PTZ-eiLYTAC and quantified HER2 levels after 48 hours using western blotting. As shown in **Figure** 4A, HER2 exhibited significant degradation. In contrast, the pertuzumab control antibody did not induce HER2 degradation (Figure S6A). Additionally, we observed degradation of EGFR, which forms a heterodimer with HER2, in JIMT-1 cells incubated with PTZ-eiLYTAC (Figure 4A). This suggests that eiLYTACs may be effective for drug-resistant tumors resulting from the activation of bypassing signaling pathways. Furthermore, we evaluated PTZ-eiLYTAC mediated HER2 degradation in nude mice with JIMT-1 xenograft tumors measuring around 100 mm^3^. Pertuzumab was included as a control. Our data demonstrated that a single intraperitoneal injection of Ptz-eiLYTAC at 30 mg/kg significantly reduced HER2 protein levels after 48 hours in the JIMT-1 tumor tissue, compared to the pertuzumab control (Figure 4B). Consistently, simultaneous degradation of EGFR was also observed in the nude mice dosed with Ptz-eiLYTAC. These findings suggest that PTZ-eiLYTAC can effectively reach the tumor site through intraperitoneal administration, leading to the degradation of HER2 homo-or heterodimers, without the need for intratumor injection as reported in other degradation approaches ^[3, 8]^. While no degradation of HER3 was observed in JIMT-1 cells, which may be attributed to the significantly lower content of HER3 in JIMT-1 cells compared to EGFR or HER2 ^[27]^, our results showed that incubation with PTZ-eiLYTAC can simultaneously degrade EGFR, HER2 and HER3 in both BT474 and T47D breast cancer cell lines (Figure 4C and 4D), indicating that Ptz-eiLYTAC may play an important therapeutic role in the drug resistance signaling pathway induced by EGFR or HER3 activation.

**Figure 4.**
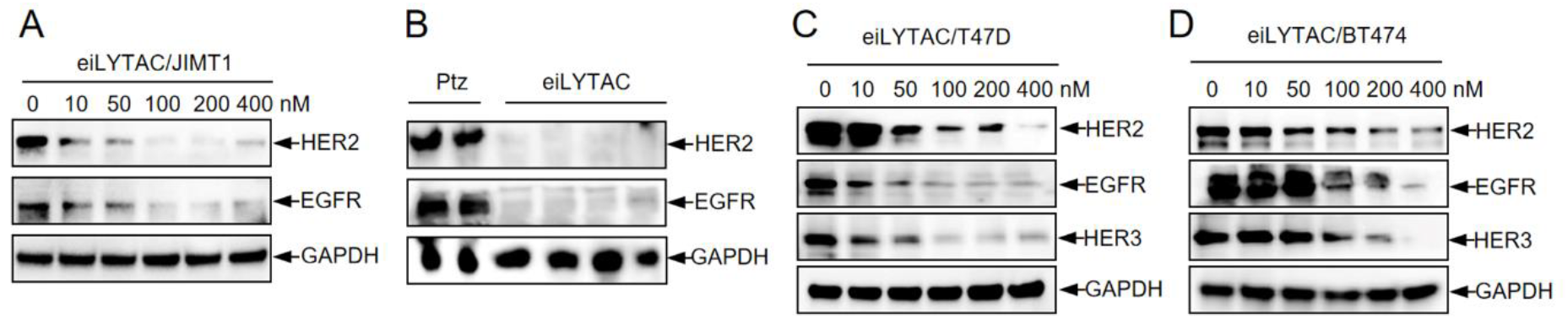
Simultaneous degradation of HER2, EGFR, and/or HER3 by Ptz-eiLYTAC both *in vitro* and *in vivo*. A) Western blot analysis showing HER2 and EGFR degradation in JIMT-1 breast cancer cells following incubation with 0 to 400 nM of Ptz-eiLYTAC for 48 h. B) Nude mice were dosed with 30 mg/kg of Ptz or Ptz-eiLYTAC. After 48 h, tumors were collected, and the expression of EGFR and HER2 was examined by western blot. C) Expression of EGFR, HER2, and HER3 was analyzed by western blot in T47D cells incubated with 0 to 400 nM of Ptz-eiLYTAC. D) Expression of EGFR, HER2, and HER3 was analyzed by western blot in BT474 cells incubated with 0-400 nM of Ptz-eiLYTAC.

Clinically, the co-occurrence of EGFR mutations and c-Met overexpression creates a challenging scenario for effective treatment. The activation of c-Met signaling pathways can bypass EGFR inhibition and promote cancer cell survival and proliferation. This phenomenon is often observed when tumors develop resistance to EGFR inhibitors, such as tyrosine kinase inhibitors (TKIs) or monoclonal antibodies ^[30]^. Thus, we incubated Cet-eiLYTAC with NCI-H1975 cells and examined EGFR and c-Met degradation using western blot analysis. As shown in Figure S7, NCI-H1975 cells incubated with the EGFR-targeting Cet-eiLYTAC exhibited simultaneous degradation of both EGFR and c-Met. Thus, eiLYTACs mediated co-degradation of its target and their binding partners should be not limited to specific targets. To our best knowledge, this is the first degrader molecular that harnessing the degradation machinery for simultaneous degradation of multiple tumor related targets.

### 2.4 EiLYTAC Inhibited Tumor Growth in Xenograft Mice Model

The efficacy of Ptz-eiLYTAC in degrading HER2 and reducing breast cancer proliferation *in vitro* prompted us to investigate its effectiveness *in vivo*. The animal experiments described were conducted at the Shenzhen Bay Laboratory, adhering to ethical regulations approved by the Administrative Panel on Laboratory Animal Care (AEWLM202201). First, we determined the *in vivo* half-life of Ptz-eiLYTAC in mouse by ELISA assays. After a single i.v. injection of 10 mg/kg Ptz-eiLYTAC, Ptz-eiLYTAC reached its peak concentration of 702.16 nM at 30 min and then gradually decreased to 194.95 nM at 480 h. The terminal half-life was 89.51 h, and the AUClast was 53064.81 nM·h as determined by Phoenix WinNonlin software (**Figure** 5A).

We then inoculated nude mice s.c. with JIMT-1 cells. After the tumor size reached about 100 mm^3^, the mice were treated twice weekly by i.p. injection of 10 mg/kg Ptz-eiLYTAC for the first 3 weeks and then once weekly for the following 1 week. PBS, pertuzumab and transtuzumab were also dosed as control groups, following the same dosing regimen as Ptz-eiLYTAC. Tumor size was measured twice per week and compared among the different groups. The results demonstrated that Ptz-eiLYTAC treatment significantly inhibited tumor growth in the mice injected with JIMT-1 cells (Figure 5B). At the end of the study, the average tumor size was 196.51 mm^3^ in the Ptz-eiLYTAC treated group, significantly smaller than the transtuzumab treated group (753.01 mm^3^), the transtuzumab treated group (742.75 mm^3^), and the PBS control group (1147.98 mm^3^). Photographic evidence of the tumors in the different treatment groups is displayed in Figure 5C. The weight of tumors in the eiLYTAC-treated groups was significantly lighter compared to the trastuzumab control group, suggesting that eiLYTAC exhibited greater potency (Figure 5D). Importantly, no toxicity or behavioral abnormalities were observed in the Ptz-eiLYTAC treated mice. Food consumption and social behaviors are similar to the control group mice. Body weights were not significantly different between Ptz-eiLYTAC-treated and vehicle control mice (Figure 5E), and no noticeable elevations of ALT or AST were observed in the mice (Figure 5F). Taken together, our *in vivo* study results indicate that the eiLYTAC is active in degrading the targeted protein and demonstrate significant anti-tumor activity *in vivo* and should be evaluated further as a candidate therapeutic for the treatment of Herceptin resistant breast cancer.

**Figure 5.**
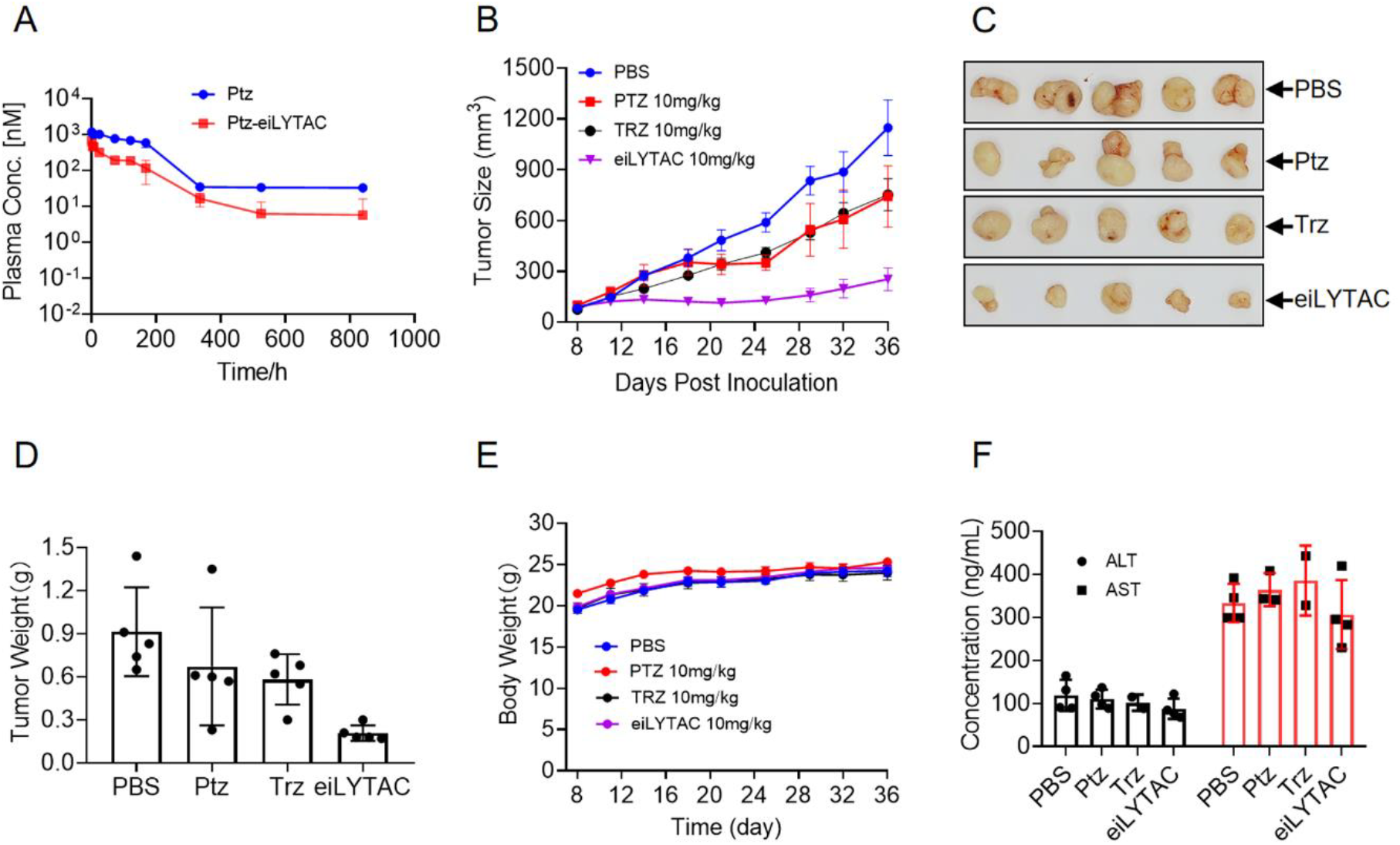
Inhibition of JIMT1 xenograft tumor growth by Ptz-eiLYTAC. A) Pharmacokinetics of Ptz and Ptz-eiLYTAC. Serum concentrations of Ptz or Ptz-eiLYTAC were measured by ELISA following a single intravenous (i.v.) injection of 10 mg/kg. B) Inhibition of JIMT-1 cells inoculated into nude mice. After the tumor size reached 100 mm^3^, Ptz or Ptz-eiLYTAC was delivered i.v. at 10 mg/kg body weight, twice a week for the first 3 weeks and once a week for the following 1 week. C) Photographs of tumor tissues dissected at day 36. D) Weight of the dissected tumor tissue. E) Measurement of body weight of mice in each group. F) Quantitative analyses of alanine aminotransferase (ALT, left panel) and aspartate aminotransferase (AST, right panel) in the serum of mice that received Ptz, Trz or Ptz-eiLYTAC treatments. Serum samples from different groups were collected at day 36, and the concentration of ALT or AST was quantified using an ELISA kit. ALT: alanine aminotransferase; AST: aspartate aminotransferase.

## 3. Conclusion

In conclusion, we have demonstrated the successful engineering of IGF2 mutant that greatly enhance its affinity toward IGF2R, while minimized its activity toward IGF1R. We have applied this mutant IGF2 to LYTAC by fusing to antibody for the development of the membrane degradation eiLYTACs that could mediate target membrane protein degradation with high efficacy both in cells and in xenograft animal models. In contrast to the recently reported LYTACs or iLYTACs technologies, eiLYTACs offer three distinct advantages. Firstly, the engineered IGF2-M5.6 selectively binds to the lysosome trafficking membrane protein IGF2R with minimal affinity for IGF1R. This significantly reduces the risk of activating IGF1R tyrosine kinase activity and the subsequent tumor formation. Secondly, the production of eiLYTACs through fusion expression in mammalian cells is straightforward. Combined with their exceptional binding affinity to IGF2R, eiLYTACs present a promising option for protein degradation and the treatment of solid tumors. Lastly, the HER2-targeting Ptz-eiLYTAC has the ability to simultaneously degrade EGFR, HER2, and/or HER3 both *in vitro* and *in vivo*. This feature is particularly advantageous for drug-resistant tumors that arise from the activation of bypass signaling pathways. As far as we know, eiLYTAC protein is the only reported tool that can achieve dual targets or triple targets degradation by targeting one membrane protein, which should provide strong support for drug development. Moreover, our eiLYTAC fusion proteins can be prepared through mammalian cell expression without chemical modification, which offers advantages in the CMC and drug development.

## 4. Experimental Section

### Cell Culture

Adherent cells were cultured in 10 cm plates at 37 °C and 5% CO2. The human cancer SK-BR-3 (Cat# CL-0211), BT474 (Cat# CL-0040), NCI-H1975 (Cat# CL-0298) were purchased from Procell Life Science & Technology Co., JIMT-1 (Cat# ZQ0950) were purchased from Cell Research Company, NCI-H1975 cells that harboring L858R/T790M/C797S EGFR mutants (Cat# CBP73213) was purchased from Nan Jing Kebai biotech Co. HEK-293F cells were purchased from Thermo Fisher Scientific company. SK-BR-3 cells were maintained in McCoy’s 5A culture medium (Procell Cat# CM-0211), BT474, T47D and NCI-H1975 cells that harboring L858R/T790M mutants were maintained in RPMI 1640 culture medium (Procell, CM-0040). JIMT-1 cells were maintained in DMEM culture medium (Gibco, Cat# 10566016). All of the culture mediums were supplemented with 10 % FBS and 1% P/S. Expi293 Expression Medium (Thermo Fisher Scientific company) was used for maintenance of cells at 37 °C, 8% CO2 with rotation (120 r.p.m.).

### Antibody Expression and Purification

In this study, DNA encoding the antibody heavy and light chains was synthesized through gene synthesis. These DNA sequences were then inserted into mammalian expression vectors called pCDNA3.4. The resulting plasmids containing the antibody genes were transfected into Expi-293F cells, which are a type of mammalian cell line commonly used for protein expression. To initiate the transfection process, the two pCDNA3.4 plasmids, each containing the gene for the light chain and heavy chain of the respective antibody, were mixed in a 1:1 ratio. Polyethylenimine (PEI) was used as a transfection reagent and added to the cell culture at a final concentration of 1 μg/mL. PEI facilitates the introduction of the plasmid DNA into the cells. After transfection, the cells were grown in a suitable culture medium for 48 hours. At this point, sodium butyrate was added to the culture medium at a final concentration of 1 mM. The cells were allowed to continue growing and producing antibodies for a total of five days after transfection. During this time, the antibodies were secreted into the cell culture supernatant. Antibodies were then purified from the supernatant using protein A affinity chromatography, a common method for isolating antibodies.

### ELISA Binding Assay

An ELISA plate from NEST (Catalog No: 520124) was used. The plate was coated overnight at 4 °C with recombinant human IGF2R or IGF1R, which were diluted to a concentration of 1 mg/L in carbonate coating buffer. The carbonate coating buffer was prepared by dissolving 1.59 g/L Na2CO3 and 2.93 g/L NaHCO_3_ in water, with a final pH value of 9.6. After coating, the wells of the ELISA plate were washed three times with 0.05% Tween 20 in PBS (PBST) to remove any unbound fractions of antibody. Subsequently, the wells were blocked with a blocking solution consisting of 2% BSA and 0.2% Tween 20 in PBS. The blocking step was carried out for 1.5 hours at 37°C. Following the blocking step, the plate was washed four times with PBST. Different concentrations of recombinant human hFc-IGF2 analogues: wild-type IGF2, M5.1 (1-156, V43M +E6R+R37A+R68A), M5.2 (1-104, V43M +E6R +R37A+R68A), M5.3 (1-67, V43M +E6R +R37A), M5.4 (1-67, V43M +E6R +R37A+V14T), M5.5 (1-67, V43M +E6R +R37A+D15A), M5.6 (1-67, V43M +E6R +R37A+F19L), M5.7 (1-67, V43M +E6R +R37A+V14T+ D15A), M5.8 (1-67, V43M +E6R +R37A+V14T +F19L), M5.9 (1-67, V43M +E6R +R37A+D15A +F19L), M5.10 (1-67, V43M +E6R +R37A+V14T+D15A +F19L) were diluted with PBS and added to the wells. The plate was then incubated on a shaker for 1 hour at 37°C to allow binding between the hFc-IGF2 mutants and the coated receptors. After incubation, the unbound fraction was washed away by washing the plate four times with PBST. An HRP-conjugated anti-human hFc secondary antibody from Jackson (Catalog No: 109035008) was diluted at a ratio of 1:5000 and added to the wells. The plate was again incubated on a shaker at 37°C for 1.5 hours to allow binding between the secondary antibody and the bound hFc-IGF2 mutants. Following this incubation, the plate was washed five times with PBST. To detect the HRP activity, a substrate for HRP was added to the wells and incubated for 5 minutes. The specific substrate used in this study was obtained from Solarbio (Catalog No: PR1200). The substrate undergoes a reaction with the HRP enzyme, resulting in a measurable signal. The reaction was then stopped by the addition of 1 M H2SO4. The optical densities of the wells were measured at 450 nm using a BioTek instrument(Synergy Neo2,S210000817), which allows for quantification of the signal generated in each well. This measurement provides information about the binding of the hFc-IGF2 mutants to the coated receptors and allows for the determination of their affinity or concentration.

### Internalization Analysis

SK-BR-3 cells were plated in the Nunc™ Lab-Tek™ II Chamber cover slide (Thermo Fisher Scientific company) 48 hours before the experiment. Cells were incubated with 200 μl of complete growth medium with 10 nM of pertuzumab or Ptz-eiLYTAC, and 20 nM of goat anti-Human, Alexa Fluor 647 for 1h. Cells were washed with HBSS for three times and incubated with 65 nM Lysotracker in HBSS for 30 min at 37 °C. Cells were then washed with HBSS three times, incubated with 1 μg/mL of Hoechst for 5 min and washed for three times before imaged by confocal microscopy.

### Protein Degradation Analysis by Western Blot

Adherent cells were plated in a 6-well plate with a concentration of 3,000,000 cells per well for 24 hours. Cells were incubated with 2 mL of complete growth medium with different concentrations of eiLYTACs or controls for 48 h. Cells were then washed with PBS for three times and lysed with RIPA buffer supplemented with protease inhibitor cocktail (Roche) on ice for 10 min. The cells were scraped, transferred to Eppendorf tubes and centrifuged at 16,000 g for 30 min at 4 °C. The supernatant was collected, and protein concentration was determined by BCA assay (Pierce). Equal amounts of lysate were loaded onto a 4–20% sodium dodecyl sulfate-polyacrylamide gel (SDS-PAGE) for electrophoresis and transferred to a 0.22 μm polyvinylidene fluoride (PVDF) membrane (Millipore). The membrane was blocked with 3 % bovine serum albumin in TBST buffer (20 mM Tris, pH 8.0, 150 mM NaCl, 0.1 % Tween-20) overnight at 4 °C and respectively incubated with primary antibodies (HER2 antibody, rabbit, 1:500; GAPDH antibody, rabbit, 1:1,000; EGFR antibody, rabbit, 1:1,000; HER3 antibody, rabbit, 1:1,000, c-Met antibody, rabbit, 1:1,000) for 1h at room temperature. After that, the membranes were washed three times with TBST buffer and then incubated with horseradish peroxidase (HRP)-conjugated secondary antibodies (1:5,000 dilution) for 1 h at room temperature. Finally, the membranes were washed five times with TBST buffer before visualization with an electro-chemiluminescence (ECL) western blotting substrate (Thermo Fisher Scientific company, Cat# A38555). For western blot analysis of Ptz-eiLYTAC mediated HER2 degradation in JIMT-1 xenograft, Five-week-old female BALB /c nude mice was used to build the JIMT-1 xenograft model. When the volume of tumors reached around 100 mm^3^ the mice were intraperitoneally injected with 30mg/kg of either pertuzumab or Ptz-eiLYTAC respectively. Mice were sacrificed and tumor tissue were homogenized for Western blot analysis. The amount of EGFR, HER2 and HER3 were determined as described previously.

### Confocal Microscopy for Membrane Protein Degradation

Adherent cells were seeded in Nunc™ Lab-Tek™ II Chamber cover slides at a density of 30,000 cells per well, 24 hours prior to the experiment. The cells were treated with 100 nM of eiLYTACs or control substances for 48 hours. The eiLYTACs or controls were added to the cells and incubated for the specified duration. Following the treatment, the cells were washed with cold PBS (phosphate-buffered saline) to remove any residual substances or media. The cells were then fixed with 4% paraformaldehyde in PBS for 15 minutes at room temperature. After fixation, the cells were washed three times with PBS to remove the fixative. The cells were then permeabilized by incubating with 0.1% Triton X-100 in PBS for 10 minutes at room temperature. After permeabilization, the cells were washed with PBST (PBS containing 0.05% Tween 20) three times to remove the permeabilization agent. The cells were then blocked with 3% BSA (bovine serum albumin) in PBST for 1 hour at room temperature. Next, the cells were incubated with a primary antibody specific to HER2/ErbB2 Rabbit mAb. The primary antibody was diluted in PBST containing 1% BSA and incubated with the cells overnight at a dilution ratio of 1:800 at room temperature for 1 hour. After the incubation with the primary antibody, the cells were washed with PBST three times for 10 minutes each at room temperature. The cells were then incubated with a secondary antibody (Donkey anti-rabbit, Alexfluor 488) diluted at a dilution ratio of 1:1000 in 1% BSA-containing PBST buffer for 40 minutes at room temperature. Following the incubation with the secondary antibody, the cells were washed three times with PBST to remove any unbound secondary antibody. To visualize the cell nuclei, the cells were incubated with 1μg/ml Hoechst dye diluted in PBST at room temperature for 10 minutes. Hoechst dye stains the cell nuclei, allowing for their visualization. Finally, the cells were imaged using a Zeiss LSM980 Confocal Laser Scanning Microscope with a 20×objective. The images obtained from each filter, including the ones for HER2/ErbB2 staining, secondary antibody staining, and Hoechst staining, were combined to produce an overlay image, allowing for the visualization and localization of HER2/ErbB2 in the cells.

### Antiproliferation Assay

Cells were seeded in a 96-well plate at a concentration of 2000 cells per well or in a black 384-well plate at a density of 800 cells per well. The plates were incubated overnight in full serum to allow the cells to attach and grow. After the initial incubation, the cells were treated with various concentrations of eiLYTACs or control antibodies at 37 °C. To measure cell proliferation, the CellTiter-Glo assay (Promega) was used. The plates were treated with CellTiter-Glo reagent, which induces cell lysis and generates a luminescent signal proportional to the amount of ATP present in the cells. In the percent proliferation calculation, cells were seeded in a 96-well plate at a concentration of 2000 cells per well. After 24 hours of incubation, the cells were treated with different concentrations of eiLYTACs or the control antibody. The cells were imaged every 6 hours for the indicated times using the IncuCyte S3 Live-Cell Analysis system. The phase-imaging channel and a ×10 objective was used to capture images of the cells. The phase confluence values, which represent the cell density or coverage in each well, were determined from the images. The confluence values were then normalized by setting the confluence of the untreated wells as the “max value,” which was equivalent to 100% proliferation. This normalization allows for the comparison of proliferation rates between different treatment conditions.

### In vivo Pharmacokinetic Study

The animal experiments described were conducted at the Shenzhen Bay Laboratory in compliance with ethical regulations approved by the Administrative Panel on Laboratory Animal Care (AEWLM202201). The laboratory ensured proper housing conditions for the animals throughout the study. SIR mice were used, with four mice assigned to each experimental group. The mice received intravenous administration of a single 10 mg/kg bolus of Ptz-eiLYTAC or pertuzumab. Blood samples were collected at specific time points, including 0.5, 3, 7, and 24 hours, as well as 3, 5, 7, 10,14,28 and 35 days post-dose. EDTA-stabilized plasma was obtained from the blood samples. The collected blood samples were processed to obtain serum by centrifugation at 4000 rpm for 30 minutes at room temperature. The serum samples were stored at -80°C until further analysis. To determine the concentrations of total antibody and eiLYTAC in the plasma samples, ELISA (Enzyme-Linked Immunosorbent Assay) was performed. A small volume (2 μl) of plasma was diluted for the ELISA assay. The ELISA standard curve was generated using the “drc” package of R software, which involved analyzing the OD450 nm values of antibody standards at different concentrations. The standard curve allowed for the determination of the “slope,” “lower,” “upper,” and “ED50” parameters. By inputting the ELISA OD450 value of the test sample into the standard curve, the blood drug concentration could be obtained. Liver biochemical parameters, specifically ALT (Abcam, ab282882) and AST (Abcam, ab263882), were analyzed using a commercially available kit. Data obtained from the experiments were analyzed using GraphPad Prism software, which is commonly used for statistical analysis and graphing in biomedical research.

### In vivo Anti-tumor Efficacy of eiLYTAC on BALB/c Nude Mice Bearing JIMT-1 Xenografts

Female BALB/c nude mice were obtained from GemPharmatech company. Five-week-old mice were selected for the study and were allowed to acclimate for one week before the experiment began. To initiate tumor growth, JIMT-1 cells were suspended in 0.2 mL of PBS containing 50% Matrigel matrix, and 1 × 107 cells were subcutaneously inoculated into the right dorsal flank of each mouse. Once the tumor volume reached approximately 100 mm^3^, the mice were randomly divided into four groups. The groups received different intraperitoneal injections: 1) PBS, 2) 10 mg/kg pertuzumab, 3) 10 mg/kg Ptz-eiLYTAC, or 4) 10 mg/kg transtuzumab. The administration frequency varied over the course of the experiment. For the first 3 weeks, the mice were dosed twice per week, while for the last one week, the dosing frequency was reduced to once per week. Tumor volume was monitored throughout the study and calculated using the formula V = (length × width^2^)/2. This formula estimates the tumor volume based on its length and width measurements. All animal experiments were conducted in compliance with the requirements of the ethics committee, ensuring the welfare and ethical treatment of the animals involved. Once the tumor volume reached approximately 1200 mm^3^, all mice were sacrificed. Tumors were collected to obtain their exact weight, which provides a quantitative measure of tumor growth and response to treatment. Additionally, serum samples were collected from the mice to perform ALT and AST ELISA assays.

## Supporting Information

**Figure S1.**
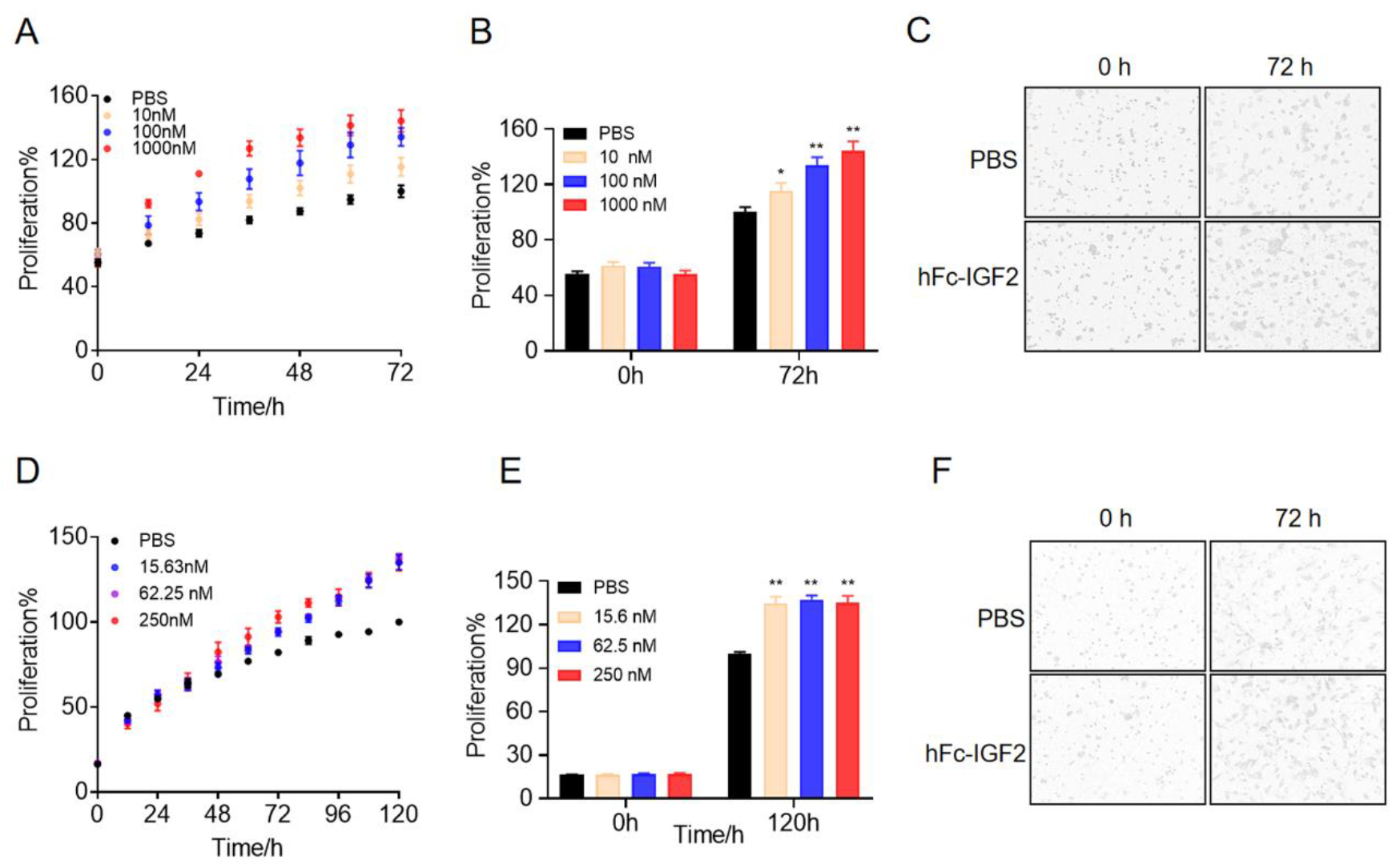
Wild-type IGF2 promotes proliferation of BT474 and SK-Hep1 cells. (A) Percentage of BT474 cell proliferation over 3 days. BT-474 cells were treated with 10 to 1000 nM of hFc-IGF2 on day 0. The proliferation percentage was measured and recorded. (B) Statistical data on cell proliferation on the first and third days. (C) Live BT474 imaging by IncuCyte throughout the 3-day period following treatment with various concentrations of hFc-IGF2. (D) to (F) Proliferation of SK-Hep1 cells was monitored by IncuCyte throughout the 5-day period following treatment with gradient concentrations of hFc-IGF2.

**Figure S2.**
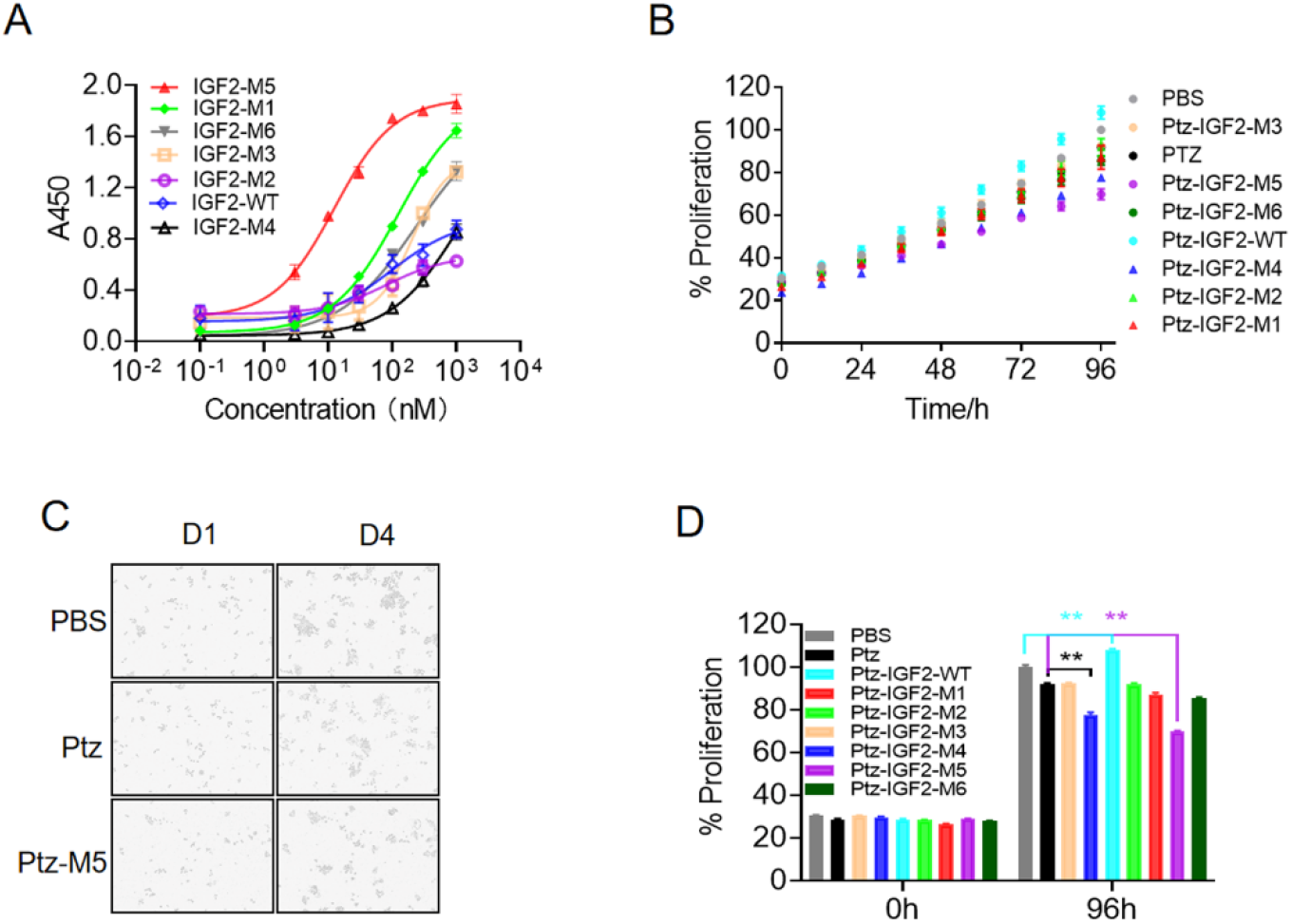
Initial screening of IGF2 mutants by ELISA and antiproliferation assay. (A) Binding affinity of different hFc-IGF2 mutants were analyzed by ELISA. The IGF2 mutants were tested for their ability to bind to domain 11 of IGF2R. (B) The proliferation of different IGF2 mutants, when fused with pertuzumab, was monitored using the IncuCyte machine. The growth and proliferation of SK-BR-3 cells in response to the different mutants were observed and analyzed. (C) Live SK-BR-3 imaging by IncuCyte at day 4 following treatment with various concentrations of hFc-IGF2. (D) The relative proliferation rate of SK-BR-3 cells was determined by comparing their growth at day1 and day4. The proliferation rates were normalized to the control group treated with PBS at day 4.

**Figure S3.**
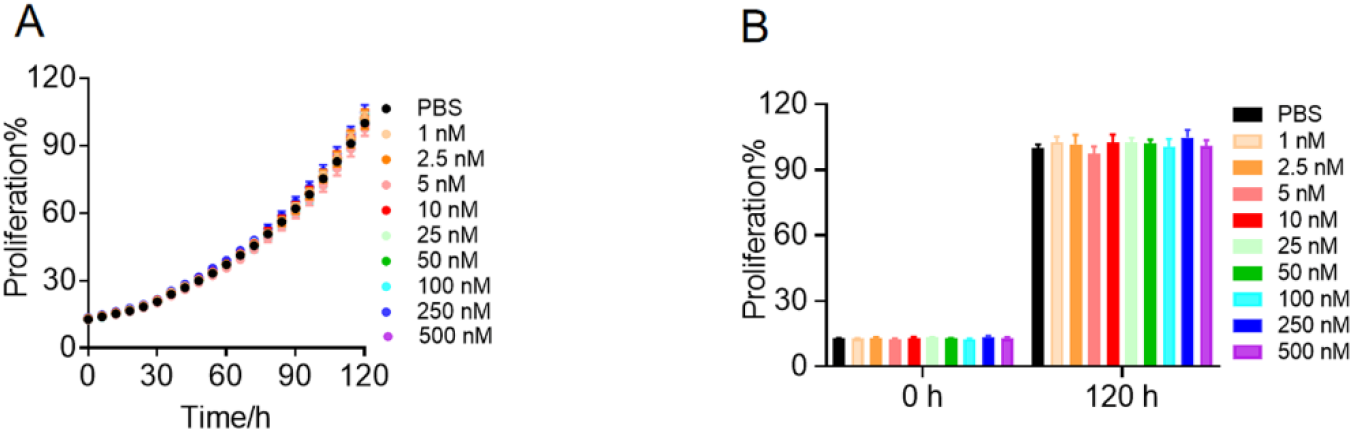
IGF2-M5.6 has no effect on proliferation of SK-BR-3 cells. (A) Percentage of SK-BR-3 cell proliferation was monitored by IncuCyte throughout the 5-day period following treatment with gradient concentrations of hFc-IGF2-M5.6. The proliferation percentage was measured and recorded. (B) Statistical data on cell proliferation on the first and fifth days.

**Figure S4.**
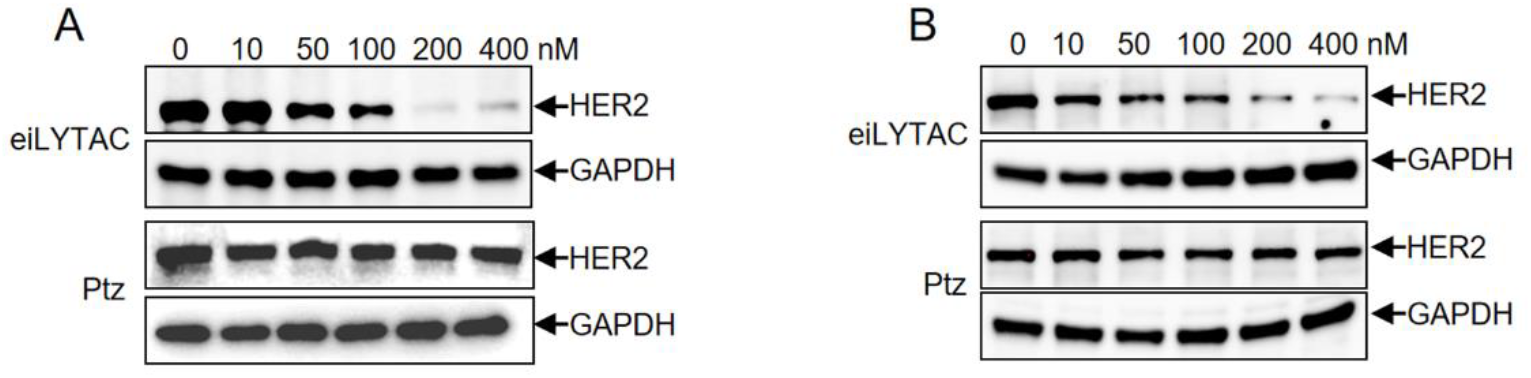
Ptz-eiLYTAC degrades HER2 in SUM159PT and MDMB435 cells. Western blot of HER2 degradation in SUM159PT (A) and MDMB435 (B) breast cancer cells, following incubation with 0-400 nM of Ptz or Ptz-eiLYTAC for 48 h.

**Figure S5.**
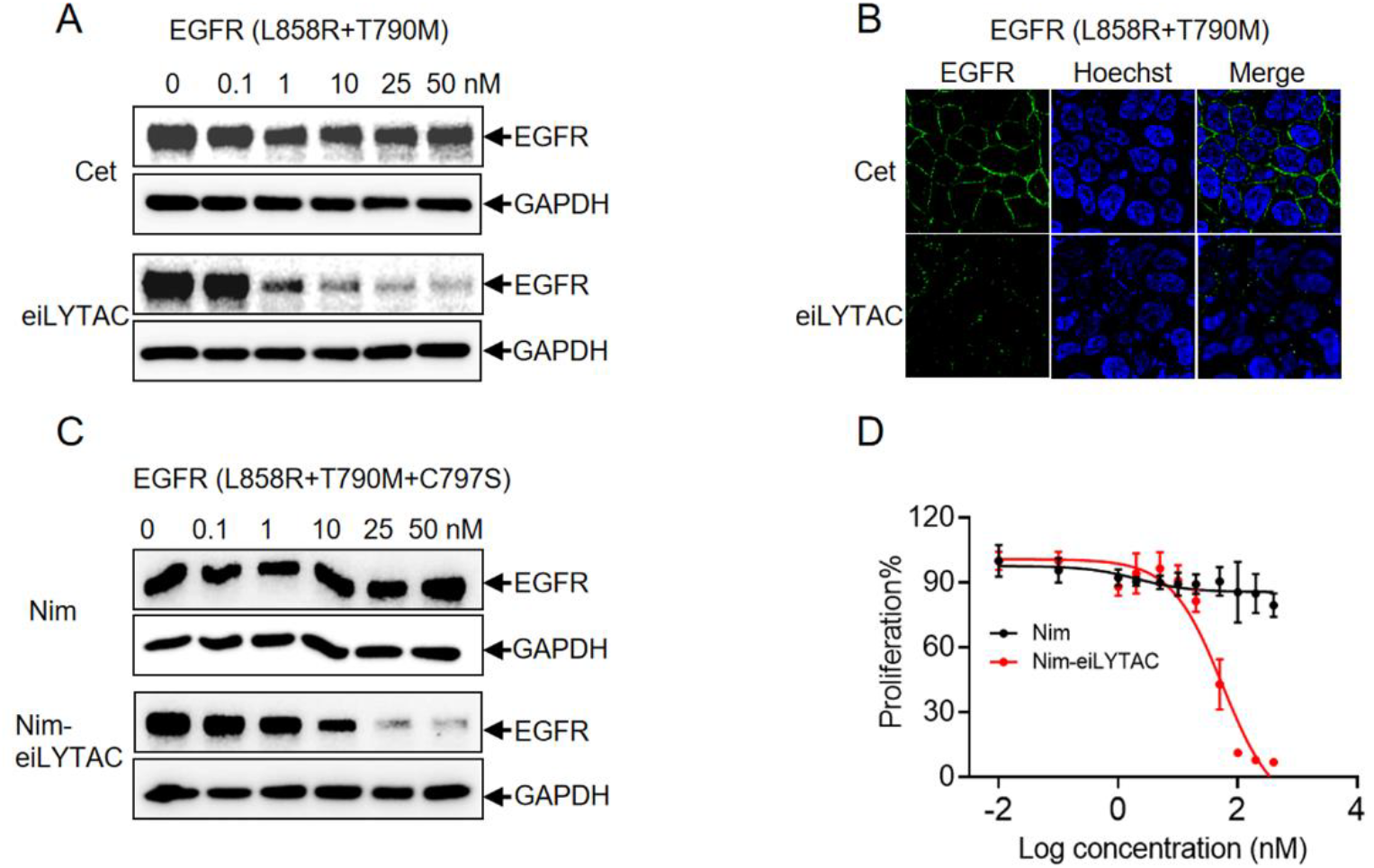
EiLYTAC degrades drug-resistant EGFR mutants and induces antiproliferation effect in non-small-cell lung cancer cells. (A) Western blot analysis of EGFR degradation in NCI-H1975 (L858R+T790M) incubated with different concentrations of Cet or Cet-eiLYTAC. (B) Confocal microscopy images showing EGFR degradation in NCI-H1975 (L858R+T790M) cells after treatment with 100 nM Cet or Cet-eiLYTAC for 48 h. (C) Western blot analysis of EGFR degradation in NCI-H1975 (L858R+T790M+C797S) NSCLC cells after incubation with 0-50 nM of Nim or Nim-eiLYTAC for 48 h. (D) Proliferation of NCI-H1975 (L858R+T790M+C797S) NSCLC cells incubated with Nim or Nim-LYTAC respectively.

**Figure S6.**
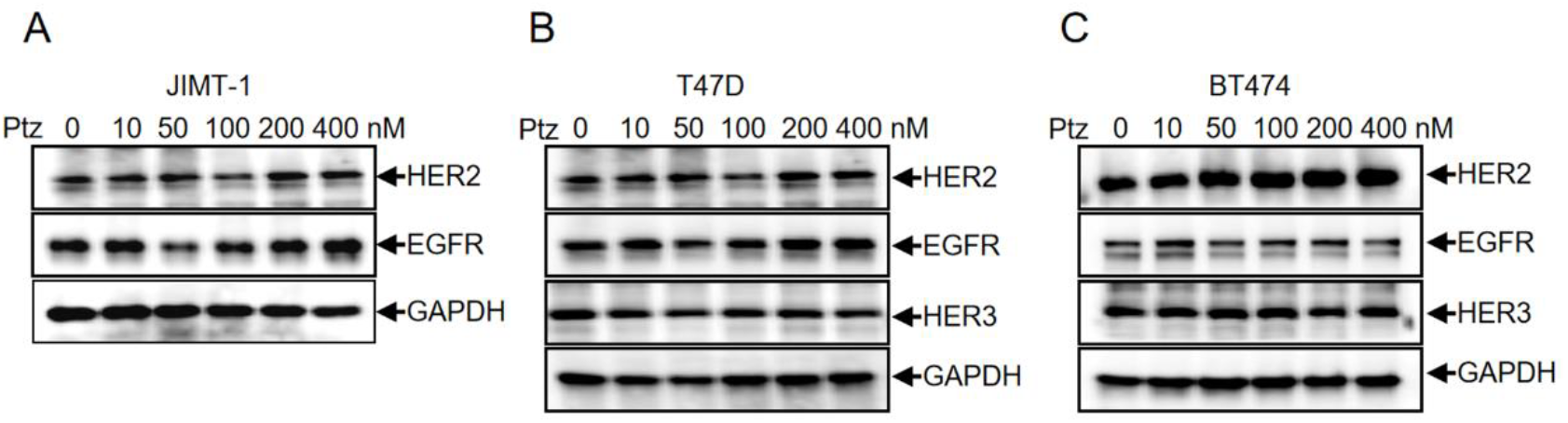
Western Blot analysis of HER2, EGFR and/or HER3 in three different breast cancer cell lines. Cells were incubated with 0-400nM of Pertuzumab for 48. After that, protein extracted in JIMT1 (A), T47D (B) and BT474 (C) was harvested and examined by Western blot.

**Figure S7.**
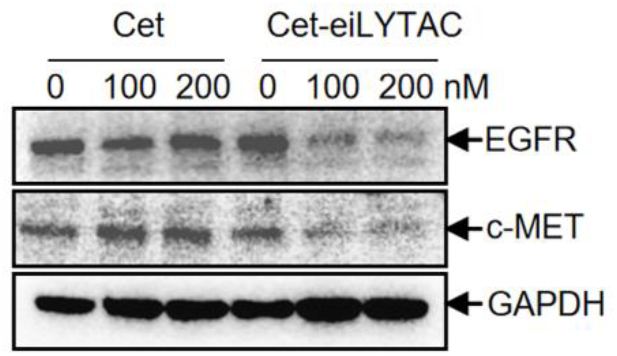
Simultaneous degradation of EGFR and c-Met by Ptz-eiLYTAC in NCI-H1975 NSCLC cells. Western blot of EGFR and c-Met degradation in NCI-H1975 cells, following incubation with 100 nM or 200 nM of Ptz-eiLYTAC for 48 h.

**Figure S8.**
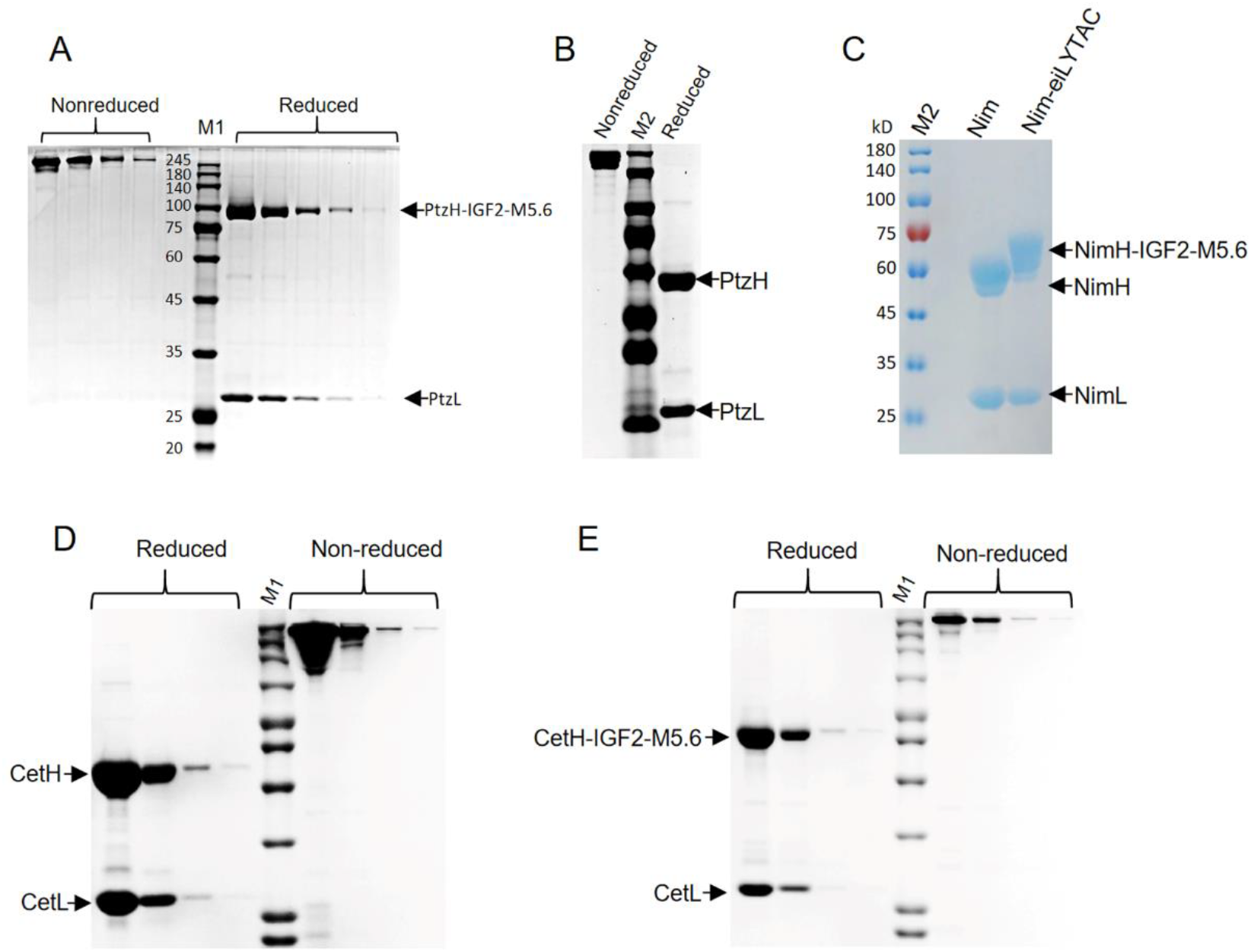
Analysis of eiLYTACs by SDSPAGE. Ptz-eiLYTAC (A), Ptz (B), Nim and Nim-eiLYTAC (C), Cet (D) and Cet-eiLYTAC (E) were expressed in HEK-293F cells and purified by Protein A-affinity chromatography. M1 and M2 are two different protein markers used.

